# SARS-CoV-2 Assembly and Egress Pathway Revealed by Correlative Multi-modal Multi-scale Cryo-imaging

**DOI:** 10.1101/2020.11.05.370239

**Authors:** Luiza Mendonça, Andrew Howe, James B. Gilchrist, Dapeng Sun, Michael L. Knight, Laura C. Zanetti-Domingues, Benji Bateman, Anna-Sophia Krebs, Long Chen, Julika Radecke, Yuewen Sheng, Vivian D. Li, Tao Ni, Ilias Kounatidis, Mohamed A. Koronfel, Marta Szynkiewicz, Maria Harkiolaki, Marisa L. Martin-Fernandez, William James, Peijun Zhang

## Abstract

Since the outbreak of the SARS-CoV-2 pandemic, there have been intense structural studies on purified recombinant viral components and inactivated viruses. However, investigation of the SARS-CoV-2 infection in the native cellular context is scarce, and there is a lack of comprehensive knowledge on SARS-CoV-2 replicative cycle. Understanding the genome replication, assembly and egress of SARS-CoV-2, a multistage process that involves different cellular compartments and the activity of many viral and cellular proteins, is critically important as it bears the means of medical intervention to stop infection. Here, we investigated SARS-CoV-2 replication in Vero cells under the near-native frozen-hydrated condition using a unique correlative multi-modal, multi-scale cryo-imaging approach combining soft X-ray cryo-tomography and serial cryoFIB/SEM volume imaging of the entire SARS-CoV-2 infected cell with cryo-electron tomography (cryoET) of cellular lamellae and cell periphery, as well as structure determination of viral components by subtomogram averaging. Our results reveal at the whole cell level profound cytopathic effects of SARS-CoV-2 infection, exemplified by a large amount of heterogeneous vesicles in the cytoplasm for RNA synthesis and virus assembly, formation of membrane tunnels through which viruses exit, and drastic cytoplasm invasion into nucleus. Furthermore, cryoET of cell lamellae reveals how viral RNAs are transported from double-membrane vesicles where they are synthesized to viral assembly sites; how viral spikes and RNPs assist in virus assembly and budding; and how fully assembled virus particles exit the cell, thus stablishing a model of SARS-CoV-2 genome replication, virus assembly and egress pathways.

## Introduction

Since December 2019, the world has been in the middle of what has been dubbed the “greatest pandemic of the century”. The etiological agent was named Severe Acute Respiratory Syndrome Coronavirus 2 (SARS-CoV-2) and the disease caused by it Coronavirus Disease 2019 (COVID-19). Coronaviruses are small enveloped viruses with positive non-segmented RNA genome. Among RNA viruses, Coronaviruses bear one of the largest genomes and its replication in the cell is complex involving frameshift slipping and replicase jumps with abundant RNA duplexes being generated. Coronaviruses, like most RNA viruses, induce the development of a range of membrane compartments that seclude and protect the viral components contributing to increased replication efficiency and innate immune recognition escape (de Wilde et al., 2013; Ertel et al., 2017; Paul et al., 2013; Snijder et al., 2020; Wolff et al., 2020b; Zhou et al., 2017).

All Coronavirus structural proteins arise from the translation of positive subgenomic RNA, which in turn are generated by replicase jumps when the negative strand copy of the viral genome is replicated. The S protein makes the viral spike, responsible for cellular attachment, entry, and fusion. It adopts two main conformations: prefusion, composed of trimers of S1 and S2, and postfusion, a non-active conformation composed solely of S2 (Cai et al., 2020; Fan et al., 2020; Hoffmann et al., 2020; Lan et al., 2020; Shang et al., 2020; Walls et al., 2020; Wang et al., 2020; Yan et al., 2020). The N protein is responsible for encapsidating and protecting the genomic viral RNA, forming ribonucleoprotein (RNP) complexes that reside in the internal space of the viral particle. The E protein is the smallest of the structural proteins and is thought to act as an ion channel (Surya et al., 2018). The M protein is the most abundant protein in SARS-CoV-2 and is a transmembrane protein that lines the internal surface of the virus lipid membrane (Neuman et al., 2011).

The coronavirus cycle starts with S interaction with ACE2 in the host cell surface (Hoffmann et al., 2020; Lan et al., 2020; Shang et al., 2020; Song et al., 2018; Wang et al., 2020; Yan et al., 2020). This interaction can either be followed by S2’ cleavage at the cell surface by TMPRSS2, or trigger the endocytosis of the viral particle, when TMPRSS2 is not present (Hoffmann et al., 2020). Upon a second still not completely characterized trigger, which may be the S2’ site cleavage (by TMPRSS2 or endosomal proteases) and/or endosomal acidification, the spike changes conformation and inserts its fusogenic peptide into the host membrane to fuse it with the viral envelope, after which the spike finally adopts the postfusion conformation (Belouzard et al., 2009; Cai et al., 2020; Fan et al., 2020; Simmons et al., 2004). The viral contents are then released into the cytoplasm, and the precursor polyproteins Pp1a and Pp1ab are synthetized. Non-structural proteins 3, 4 and 6, which are part of Pp1a/Pp1ab, induce the formation of secluded, often interconnected, membranous compartments known as DMVs (Double Membrane Vesicles) (Doyle et al., 2018; Hagemeijer et al., 2014; Oudshoorn et al., 2017). The DMVs compartmentalize the Replication Transcription Complexes (RTCs) and are the sole compartments where viral genome replication takes place (Snijder et al., 2020), both for the synthesis of the negative strand viral RNA and for the synthesis of the positive strand subgenomic mRNAs and viral genome RNA copies. Initially, it was thought that these compartments were sealed and had no connection to the cytoplasm, raising the mystery of how the mRNAs could reach the cytoplasm to be translated by the cellular ribosomes. Recently, however, a molecular pore has been described in MHV and SARS-CoV-2 that can serve as export portal for the mRNA and positive strand viral genome copies (Wolff et al., 2020a). The assembly of the viral particle is thought to take place at modified cellular membranes derived from the ER, Golgi and ERGIC (endoplasmic-reticulum-Golgi intermediate compartment), and viral release through exocytosis, based on studies of other coronaviruses (Goldsmith et al., 2004; Knoops et al., 2008; Ogando et al., 2020; Stertz et al., 2007). Although, there have been intense structural studies on recombinant viral components and purified inactivated viruses (Hoffmann et al., 2020; Ke et al., 2020; Lan et al., 2020; Shang et al., 2020; Song et al., 2018; Turoňová et al., 2020; Wang et al., 2020; Yan et al., 2020; Yao et al., 2020), investigation of the SARS-CoV-2 replication process in the native cellular context is scarce, and viral assembly and egress are still not well understood at the molecular level.

In this study we investigated SARS-CoV-2 replication in Vero cells under near-native conditions exploiting a unique correlative multi-modal multi-scale cryo-imaging approach by combining soft X-ray cryo-tomography and serial cryoFIB/SEM volume imaging of the entire SARS-CoV-2 infected cell with cryo-electron tomography (cryoET) of cell lamellae and cell periphery, as well as structure determination of viral components through subtomogram averaging. This approach empowers a holistic view of SARS-CoV-2 infection, from the whole cell to individual molecules, revealing novel pathways of SARS-CoV-2 assembly and egress and cytopathic effects of SARS-CoV-2 infection.

## Results

### SARS-CoV-2 replication induces profound cytopathic effects in host cells

To image and investigate SARS-CoV-2 replication in near-native cell context, we infected Vero cells grown on indexed EM grids with SARS-CoV-2 at 0, 0.1 and 0.5 multiplicity of infection (MOI). At 24 hours post infection (hpi), the cells were fixed with 4% paraformaldehyde and plunge frozen in liquid ethane. As illustrated in the workflow (Figure S1), cryoEM grids containing SARS-CoV-2 infected cells were first screened in a Titan Krios to identify each individual infected cell (39.2 % for MOI of 0.1 and 45.4% for MOI 0.5) where cryoET tilt series were collected at the cell periphery. The grids were then transferred to a FIB/SEM dualbeam instrument and the same infected cells were subjected to either serial cryoFIB/SEM volume imaging (Zhu et al., 2021) or cryoFIB milling of cellular lamellae where additional cryoET tilt series were collected (Sutton et al., 2020). Alternatively, we imaged infected cells on cryoEM grids by soft X-ray cryo-tomography (Harkiolaki et al., 2018; Kounatidis et al., 2020). It’s worth mentioning that serial cryoFIB/SEM volume imaging is emerging as a new cryo-volume imaging technique for the study of large volumes of near-native, fully hydrated frozen cells and tissues at voxel sizes of 10 nm and below, adding to the capability of soft X-ray cryo-tomography. These imaging modalities provide structural information at different length scales and are highly complementary. Such a unique approach enabled the comprehensive investigation of the SARS-CoV-2 replication and cytopathology effects in a multi-modal, multi-scale and correlative manner.

Compared to uninfected cells (Figure S2, Movie 1), serial cryoFIB/SEM images of SARS-CoV-2 infected cells display an extensive array of cytopathological alterations throughout the entire cell, as illustrated in Figure 1 and Movies 2-5. At the cell surface, there were many virus-containing membrane tunnels extending deep into the cell (Figure 1A, “T”, Movie 2), in addition to electron lucent membrane vesicles (Figure 1A, “V”). Virus particles were also found within intracellular vesicles not connected to cell membrane (Figure 1A, red arrow). Deep into the cell, much of the cytoplasm is occupied with abundant membrane compartments of different morphologies, including numerous vesicles (“V”) (Movie 4), the complex membrane compartment (pink arrow), the endoplasmic reticulum (ER) and the nucleus (“Nuc”) (Figure 1B, Movie 2 and 3). Most of these vesicles are the so-called “double membrane vesicles” (DMVs) where viral genome replication takes place (Snijder et al., 2020). Interestingly, we observed a different type of electron lucent vesicles which appears lined up with a string of very small vesicles (Figure 1B, “V*”), the function of which is revealed by cryoET of cell lamella discussed in sections below. At the mid-cell where the nucleus is present, cryoFIB/SEM images clearly display nuclear pores in frozen-hydrated cells (Figure 1C, Figure S2A&C, blue arrows). Mitochondria are frequently disrupted in the infected cells (Figure 1C, yellow arrow, Movie 2) compared to the uninfected cells (Figure S2, Movie 1). In addition, cryoFIB/SEM reveals electron-dense complex membrane compartments often seen in infected cells (Figure 1B, pink arrows). The more striking feature, observed in 2 of 3 infected cells imaged, is the cytopathic damage to the nucleus compared to the control cells, where nearly a half of the nucleus has been taken up by the invaginated cytoplasm (Figure 1D, Movie 5). We noticed that in a recent study, such cytoplasm invagination was also seen in one of the conventional EM images of stained plastic sections of SARS-CoV-2 infected cells, although no description was given (Lamers et al., 2020).

**Figure 1.**
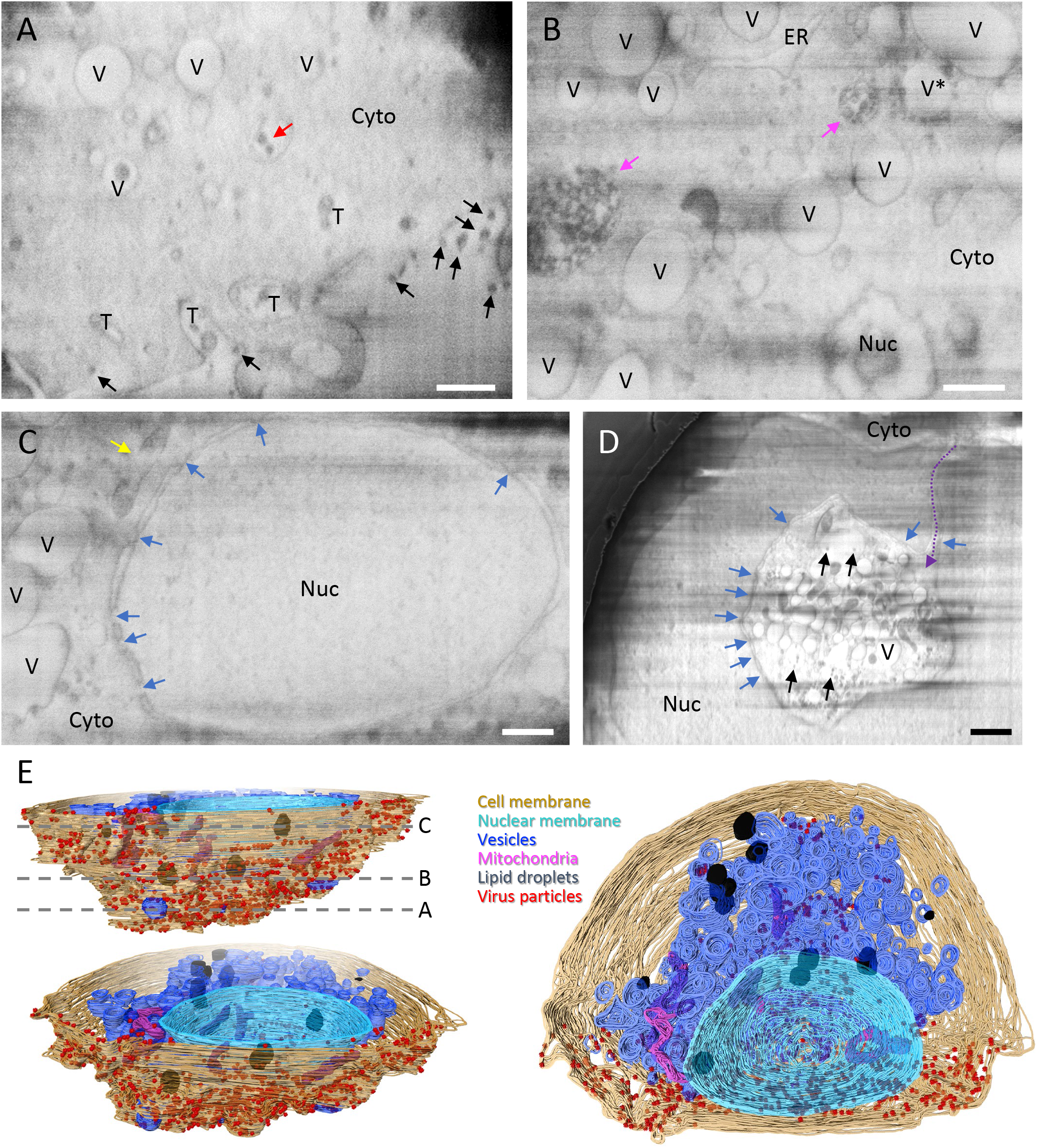
Serial cryoFIB/SEM volume imaging of entire SARS-CoV-2 infected cell. (A-D) Representative cryoFIB/SEM slices of a SARS-CoV-2 infected cell at the cell periphery (A), cytoplasm (B), cell nucleus (C), and invagination of cytoplasm into the nuclear space (note, from a different cell) (D). Scale bars, 500 nm in A-C, 1 μm in D. Black and red arrows, extracellular and intracellular virus particles; blue arrows, nuclear pores; yellow arrow, a damaged mitochondria; pink arrows, complex membrane compartment; dashed purple arrow, invagination path; V and V*, vesicles; T, tunnels; Nuc, nucleus; Cyto, cytoplasm; ER, Endoplasmic reticulum. (E) Surface rendering of the segmented volume of SARS-CoV-2 infected cell shown in A-C. Segmented organelles and virus particles are labeled with colors indicated. The dashed lines (E, top left panel) indicate the positions of slices shown in A-C, respectively.

Independently, we investigated cytopathic effect of SARS-CoV-2 infection using soft X-ray cryo-tomography. Consistent with the cryoFIB/SEM volume imaging results, the overview images of soft X-ray display profound changes in mitochondria morphology, as they appear fragmented in the SARS-CoV-2 infected cell (Figure S3B&D, F), also shown in cryoET (Figure S4), compared to the extended and networked mitochondria in the uninfected cell (Figure S3A&C). Consistent with this, SARS-CoV-2 infection leads to profound downregulation of transcriptional pathways related to mitochondrial function in human induced pluripotent stem-cell derived cardiomyocytes (Sharma et al., 2020), and cardiac complications are more common in COVID-19 patients (Fried et al., 2020). Virus particles on the cell surface are clearly distinguishable in soft X-ray cryo-tomogram (Figure S3E, black arrow). Also clearly observed are extensive vesiculation (Figure S3F) and a partial nucleus invagination in the infected cell (Figure 3G).

**Figure 2.**
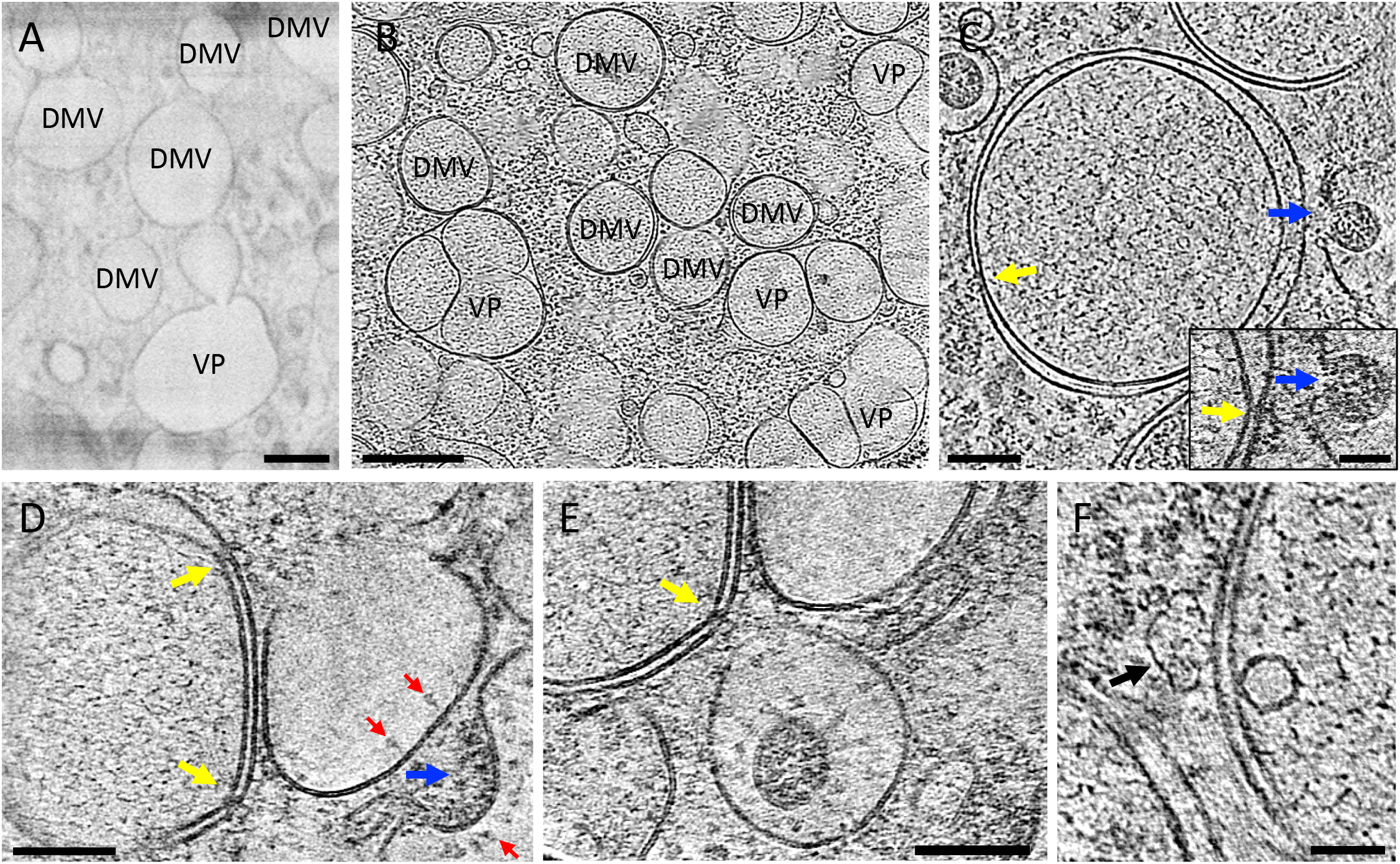
SARS-CoV-2 genome replication and RNA synthesis. (A-B) Overview of a SARS-CoV-2 infected cell in cryoFIB/SEM slice (A) and cryo-lamella tomogram slice (B) depicting double membrane vesicles (DMV) and vesicle packets (VP). (C) Cryo-tomogram slice of DMV at high magnification. Inset depicts detail of a DMV pore next to a viral assembly site. (D-E) Pores on DMVs next to assembly sites. (F) Vaultosome in close proximity to a DMV outer membrane. DMV – Double membrane vesicle. VP – Vesicle packet. Yellow arrows – DMV portals. Blue arrows – Viral assembly sites. Red arrows – Viral spikes. Black arrow – Vaultosome. Scale bars are 300 nm in A, 500 nm in B, 100 nm in C, 50 nm in C inset, 100 nm in D and E, 50 nm in F.

**Figure 3.**
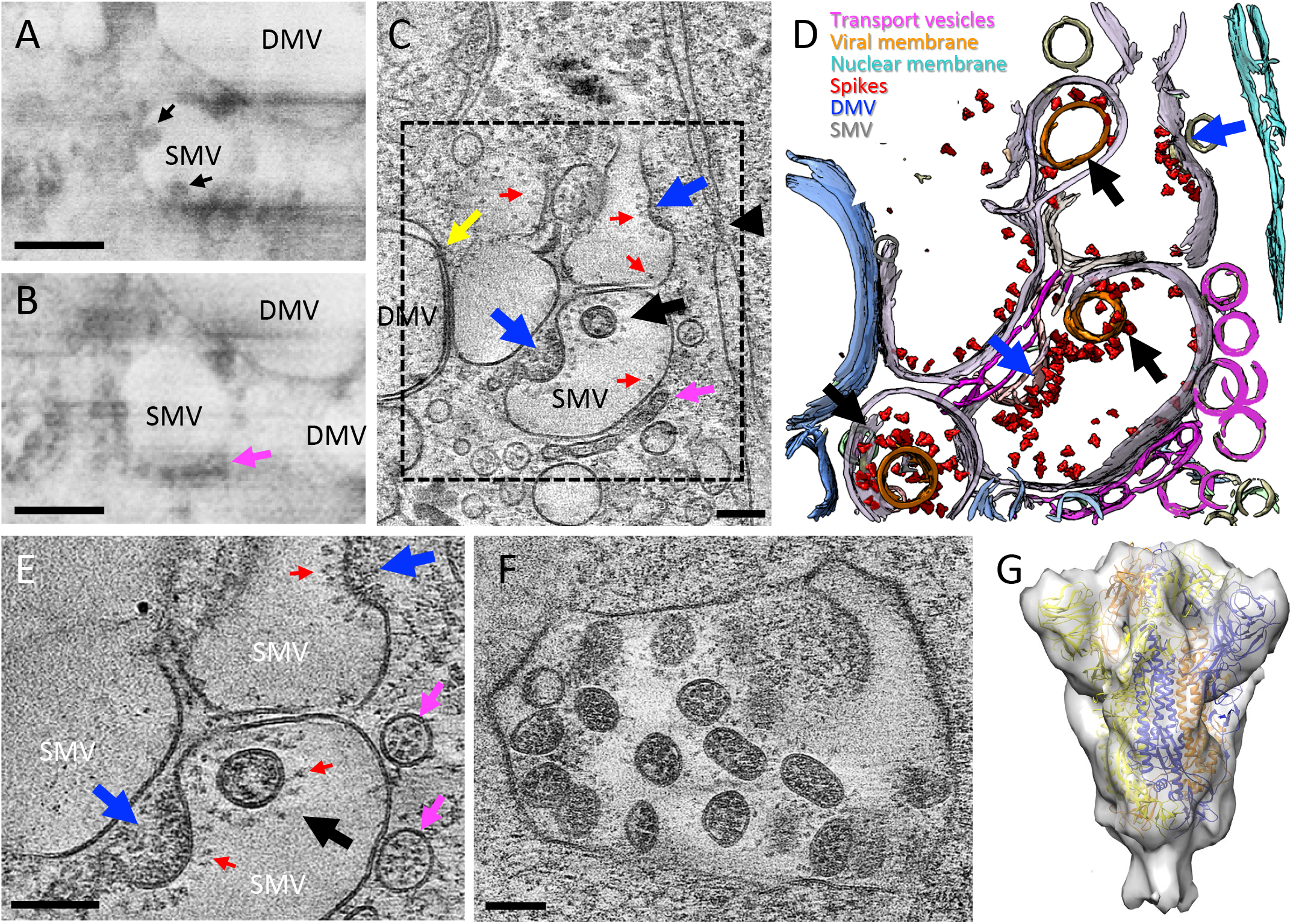
SARS-CoV-2 cytoplasmic viral assembly. (A-B) CryoFIB/SEM images of two sequential slices separated by 80 nm. Black arrows point to virus particles in single membrane vesicle (SMV). Pink arrow points to small dense vesicles lining the outside of virus-containing SMV. (C) Tomographic slice of cryoFIB lamella depicting SARS-CoV-2 assembly, with DMV portals (yellow arrow), assembling viruses (blue arrow), assembled virus (black arrow), viral spikes on SMV membranes (red arrows), dense vesicles around the assembly site (pink arrow, as in B) and a nucleopore (black arrowhead). (D) Density segmentation of C, displaying three virus particles (black arrows) and two assembly sites (blue arrows). (E) An enlarged view (at a different angle) of boxed area in C, showing assembled virus (black arrow), assembling viruses (blue arrows), spikes (red arrows) and spike-containing vesicles (pink arrows). (F) Large intracellular virus-containing vesicle (LVCV) full of readily assembled viruses. (G) Subtomogram average of viral spikes of intracellular viruses from cell lamellae at 11 Å resolution, fitted with an atomic model of spike trimer (PDB 6ZB5) (Toelzer et al., 2020). Scale bar is 300 nm in A and B; and 100 nm in C, E and F.

Altogether, soft X-ray cryo-tomography and serial cryoFIB/SEM volume imaging provide a comprehensive overview of cytopathic effects of SARS-CoV-2 infection in native cells, including membrane tunnels at the cell surface, virus-containing vesicles, intracellular complex membrane compartments, and numerous heterogeneous vesicles, invagination of nuclear membrane and damaged mitochondria, all of which can be correlated with in-depth *in cellulo* cryoET analysis (sections below).

### SARS-CoV-2 RNA synthesis and transport

The first step in SARS-CoV-2 production is viral genome replication. Coronaviruses have evolved a sophisticated RNA replication strategy for the generation of the subgenomic RNAs, which relies heavily on double stranded RNA intermediaries, a potent activator of RIG-I and MDA-5 (Andrejeva et al., 2004; Hornung et al., 2006; Neufeldt et al., 2016). Thus, cellular compartmentalization of RNA transcripts serves as an innate immune evasion strategy. DMVs are induced during the replication of a variety of RNA viruses (de Wilde et al., 2013; Ertel et al., 2017; Paul et al., 2013; Wolff et al., 2020b; Zhou et al., 2017), and were identified as the sole compartment where viral RNA transcription occurs for coronaviruses (Snijder et al., 2020). Indeed, cryoET of cell lamella revealed that abundant intracellular vesicles observed in the 3D volume of infected cell (Figure 1D, Movie 3 and 4) are clearly DMVs containing filamentous structures that likely correspond to viral RNA transcripts as previously suggested (Reggiori et al., 2010; Snijder et al., 2020) (Figure 2A-C, Movie 6). There are also a substantial amount of so-called vesicle packets (VPs) (Figure 2B) (Ogando et al., 2020), apparently resulting from the fusion of the outer membranes of DMVs. Since the sample is 24 hours post infection, this is consistent with a previous observation that the number of VPs increases with the time of infection (Snijder et al., 2020). Until very recently, DMVs were thought to be completely enclosed, which raised the question of how the viral mRNAs could gain access to the cytoplasm to be translated. We observed several double-membrane-spanning pore complexes in DMVs (Figure 2C-F, yellow arrow), resembling the RNA transport portal seen in DMVs of murine hepatitis coronavirus (MHV) infected cells in a recent study (Wolff et al., 2020a). However, the portal appears rare in DMVs of SARS-CoV-2 (total 9 portals from 24 DMVs) compared to those of MHV (8 portals per DMV), signifying the difference between coronaviruses. We also observed vaultosomes near the DMV outer membranes (Figure 2F, black arrow). It is still unclear what is the physiological role of vaults is, but they have been associated with RNA nucleocytoplasmic transport, innate immunity, and cellular stress response (Berger et al., 2009; Woodward et al., 2015), which are likely to be activated by infection.

### SARS-CoV-2 assembly and budding

The translation of the subgenomic vRNAs gives rise, amongst others, to the structural proteins N, M, E and S, which are required for assembly. M, E and S are membrane-associated proteins and are localized to the ER, Golgi and the ERGIC (Reggiori et al., 2010; Stertz et al., 2007). The N protein associates with the genomic vRNA and M protein, which presumably drives vRNA packaging and genome encapsidation (Lu et al., 2020; Vennema et al., 1996). The main assembly and budding site of other coronaviruses has been previously described at the ERGIC by conventional EM of stained, plastic embedded, and sectioned samples (de Wilde et al., 2013; Knoops et al., 2008; Reggiori et al., 2010; Stertz et al., 2007). In serial cryoFIB/SEM images of SARS-CoV-2 infected cells, we clearly observed vesicles containing virus particles (Figure 3A, black arrows), along with a string of small dense vesicles lining along the vesicle membrane in the close proximity to the DMVs (Figure 3B, pink arrow). High-resolution cryoET of cell lamella from a similar cellular region allowed us to unambiguously identify that these are in fact SARS-CoV-2 assembly and budding sites (Figure 3C-E). CryoET further revealed that the string of small dense vesicles are SARS-CoV-2 spike containing trans-Golgi transport vesicles, supplying newly synthesized spikes to the assembly sites via fusion with the SMVs (Figure 3C-E pink arrows, Movie 7). Indeed, spikes are observed on SMV membranes clustered at the assembly sites, in addition to being sparsely distributed (Figure 2D, Figure 3C-E, red arrows, Movie 7). Interestingly, several SARS-CoV-2 assembly intermediates were observed within a single tomogram from a cell lamella (Figure 3C-E, blue arrows, Movie 7), along with fully assembled virus particles released into SMVs (Figure 3C-E, black arrows, Movie 7), thus allows capturing the active assembly and budding process of SARS-CoV-2. It is conceivable that upon fusion of transport vesicles with the SMV, spikes are readily diffused onto the SMV membrane, then clustering when interacting with N-associated vRNA, possibly via M protein (Lu et al., 2020; Neuman et al., 2011), which initiates the budding process that finally releases the viral particle into the SMV. Consistent with this, spike clusters are observed exclusively associated with the agglutination/gathering of electron dense material, which presumably represents viral genome. Noticeably, the virus assembly site is frequently present in the vicinity of RNA portals in DMVs (Figure 2C-E, yellow arrows), potentially facilitating the assembly process. The N-associated vRNA further matures as distinct RNPs recognizable in the fully assembled virus particles (Movie 7).

Most virus particles are found in ERGIC SMVs, some contain a single virion, while others encompass multiple virions (Figure 3E-F). CryoET and subtomogram averaging of 450 spikes from these particles yielded a density map at 11 Å resolution (at 0.143 FSC cut-off) by emClarity (Himes and Zhang, 2018) (Figure 3F-G, Figure S5). The averaged density map resolves the overall spike structure, which overlaps very well with prefusion spike atomic models (Hoffmann et al., 2020; Lan et al., 2020; Shang et al., 2020; Walls et al., 2020; Wang et al., 2020; Yan et al., 2020). Virus particles were also observed in electron-dense complex autophagolysosome-like compartments (ALC) (Figure 4), which likely correspond to the complex membrane compartments seen in cryoFIB/SEM (Figure 1B, pink arrows). These virus particles, however, have either no spikes (Figure 4B) or a few postfusion spikes on their surfaces (Figure 4E-F). Viruses protected by single membrane vesicles (SMV) in ALC show prefusion spikes (Figure 4G-H). These could be off-pathway sites of viral assembly (in the case of SMV-protected viruses displaying prefusion spikes), remnants of late endosomes from viral entry or lysosomes for viral degradation (in the case of viruses displaying postfusion spikes or no spike). The fact that the spike proteins are in the postfusion state suggests that proteolytic processing has taken place in these compartments resulting in S1 shedding. Therefore, we suggest that assembly at the ERGIC SMVs is the only pathway which will lead to infectious viral progeny.

**Figure 4.**
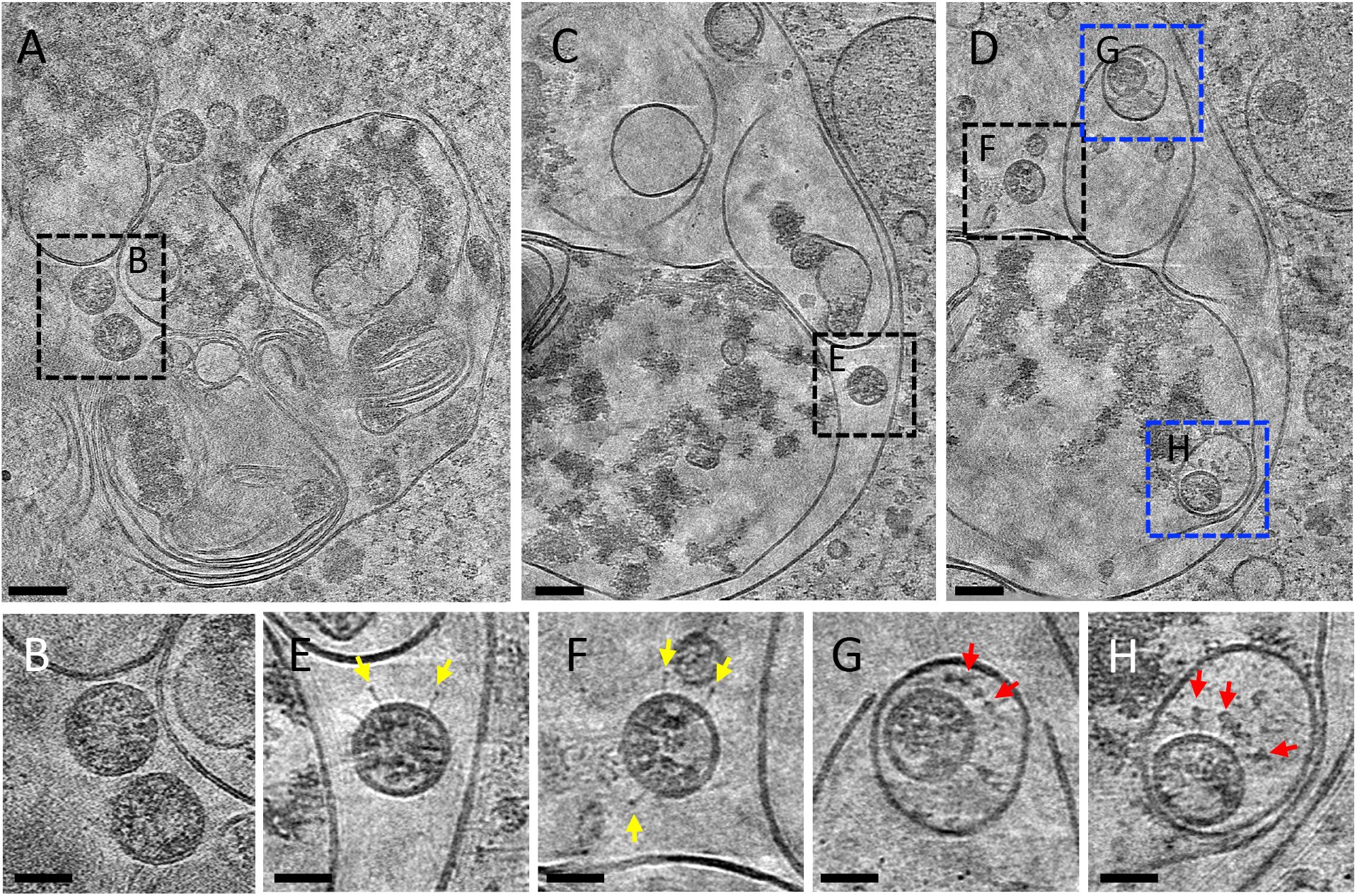
Non-productive autophagolysosome-like compartments (ALC). (A) Tomographic slice of an ALC in cell lamella depicting convoluted membranes containing virus particles (boxed area). (B) Detailed view of spikeless viruses from the boxed area in A. (C-D) Consecutive tomographic slices of the same ALC separated by 140 nm, containing viruses (black and blue boxed areas). (E-F) Detailed view of viruses with a few postfusion spikes (yellow arrows) from boxed areas in C and D. (G-H) Detailed view of viruses protected by single membrane vesicles (SMV) harboring prefusion spikes (red arrows) from blue boxed areas in D. Scale bars are 500 nm in A; 100 nm in B, C; 50 nm in C, D, E, F, G, H.

### SARS-CoV-2 egress with two distinct pathways

There has not been much detailed studies on how SARS-CoV-2 viruses are released from cell. We investigated SARS-CoV-2 egress using both serial cryoFIB/SEM volume imaging and cryoET. CryoFIB/SEM images clearly reveal virus exiting tunnels in 3D at the cell periphery connecting to cell membrane (Figure 5A-B, Movie 2). This likely resulted from the fusion of very large multi-virus containing vesicles with cell membrane, i.e. egress through exocytosis. Consistent with cryoFIB/SEM analysis, we also observed virus exiting tunnels in cryo-tomograms (Figure 5C). The fact that these compartments often contained many viral particles suggests that this is a snapshot of viral exit, rather than cellular entry.

**Figure 5.**
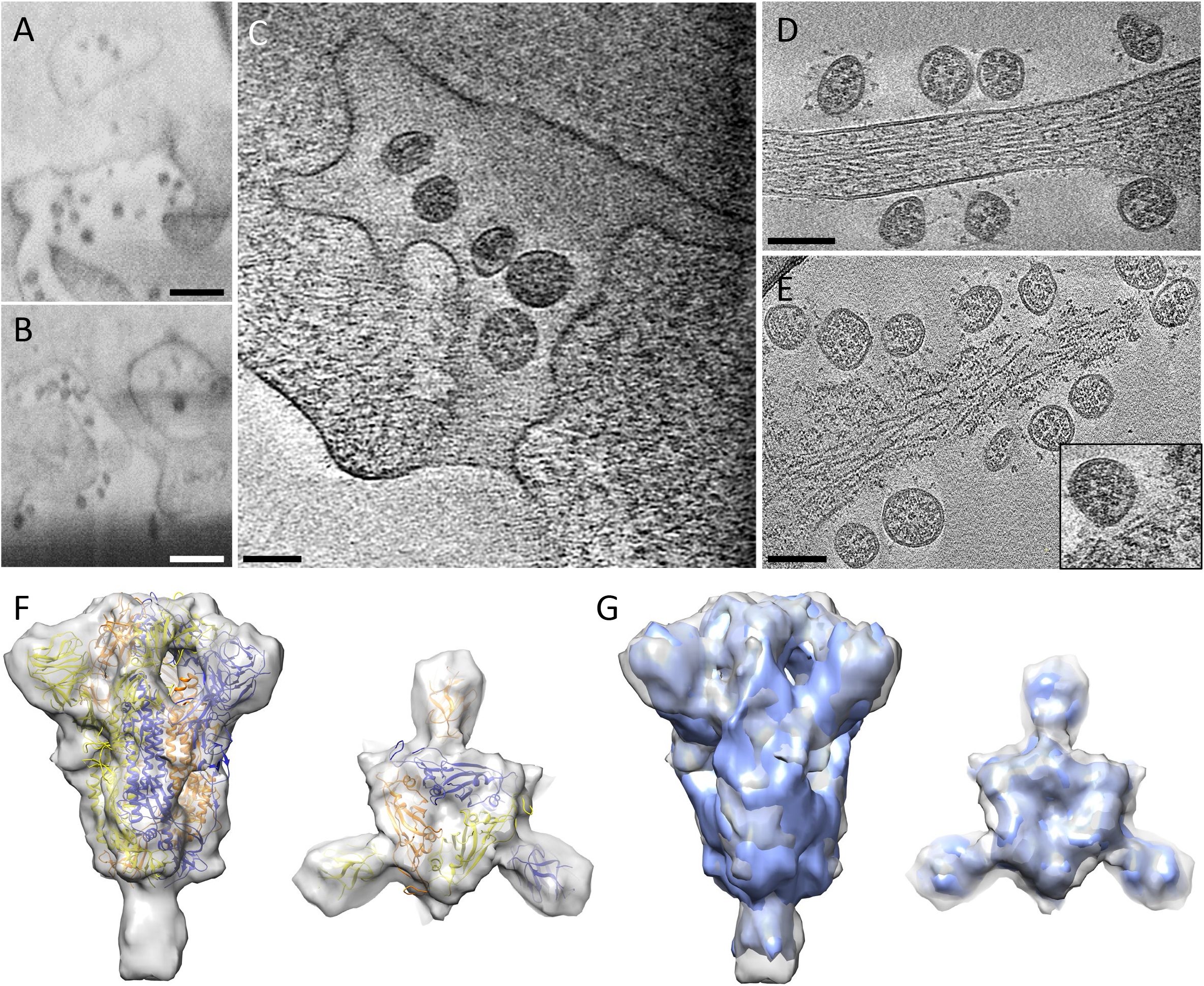
SARS-CoV-2 viral egress pathways. (A-B) CryoFIB/SEM images of cell periphery, depicting virus particles exiting through extended tunnels connected to external of the cell. (C) CryoET of the SARS-CoV-2 exiting tunnel. (D) Viruses outside of the cell. (E) Membrane-rupture viral egress. Inset, close-up views of membrane rupture sites. (F) Subtomogram average of spikes on released viruses at 9 Å resolution fitted with an atomic model of spike trimer (PDB 6ZB5) (Toelzer et al., 2020), viewed from side and top. (G) Comparison of spike structures from intracellular assembled viruses (blue) and extracellular released viruses (transparent grey), shown in side and top views. Scale bar is 300 nm in A and B, and 100 nm in C, D and E.

However, in addition to exocytosis, we also frequently found plasma membrane discontinuities next to viral particles outside the cell (Figure 5E). There are 116 membrane lesion sites next to virus particles in 74 tomograms, and 44.6% of tomograms show cell membrane lesions in infected cells (Figure 5D-E, Movie 8-9), whereas 18.7 % tomograms from uninfected cells display similar membrane lesions (10 membrane lesion sites from 16 tomograms). Close inspection of individual membrane lesions indicats that the underlying cytoskeleton, such as actin filaments, is largely intact, and only the plasma membrane was compromised (Figure 5E inset). The fact that we observed the same membrane lesion, but to a lesser extent in control cells, suggests that SARS-CoV-2 exploits the host cell machinery for its egress. Cellular exit through plasma membrane rupture may be an alternative viral egress pathway of SARS-CoV-2, as this is observed frequently in our data. It is unclear whether the cell can recover from such membrane wounds, or if exit through membrane rupture is a sign of late infection and will eventually lead to cell lysis and death. Severe cases of COVID-19 are characterized by high levels of inflammation markers (Lucas et al., 2020), and viral lysis may be one of the mechanisms triggering this response.

CryoET subtomogram averaging of 7090 spikes from extracellular virus particles yielded a density map at 8.7 Å resolution (at 0.143 FSC cut-off), which represents the prefusion state (Figure 4F, Figure S5). Spike structures from intracellular and extracellular viruses agree with each other very well (Figure 4G), suggesting that no further structure rearrangement takes place for viral spikes from assembly to egress. While all previous spike structures are either from recombinant proteins or from purified inactivated virus particles (Cai et al., 2020; Fan et al., 2020; Hoffmann et al., 2020; Ke et al., 2020; Lan et al., 2020; Shang et al., 2020; Turoňová et al., 2020; Walls et al., 2020; Wang et al., 2020; Yan et al., 2020; Yao et al., 2020), the two spike structures presented here are derived directly from infected cells in the cellular context, and thus represent the closest to native condition, providing a strong validation for these *in vitro* structures.

## Discussion

We used a multi-modal, multi-scale and correlative approach to investigate SARS-CoV-2 infection process in native cells, from the whole cell to subcellular and to the molecular level. The integration of multi-scale imaging data, achieved through this advanced workflow (Figure S1), has led us to propose a pathway for SARS-CoV-2 replication, in particular virus genome replication, assembly and egress. The replication of SARS-CoV-2 appears spatially well-organized and highly efficient. From genome replication, to protein synthesis and transport, to virus assembly and budding, these processes take place in close-knit cytoplasmic compartments. As illustrated in Figure 6, RNA replication, including vRNA and mRNA, occurs in DMVs, secluding them from innate immune response (step 1). The newly synthesized vRNAs are then transported out of DMVs through the transmembrane portals to ERGIC virus assembly sites proximal to DMVs and portals (step 2a), whereas mRNAs exit through the same portal to cytoplasm and ER/Golgi for protein production (step 2b). The viral spikes, in a trimeric prefusion form produced and matured in ER/Golgi network, are transported to the ERGIC assembly sites via small transport vesicles (step 3). Upon fusion of transport vesicles with ERGIC membranes (or SMVs), viral spikes cluster at the assembly site where vRNA and N protein are present, resulting in a positive membrane curvature and finally bud into the SMV (step 4). Depending on the number of virus particles within SMVs, there might be two distinct egress pathways: the large virus-containing vesicle (LVCV) through tunnels via exocytosis (step 5a), and the single virus-containing vesicle (SVCV) breaking out through cell membrane rupture (step 5b), although it is still unclear what is the mechanism of membrane rupture exploited by the virus.

**Figure 6.**
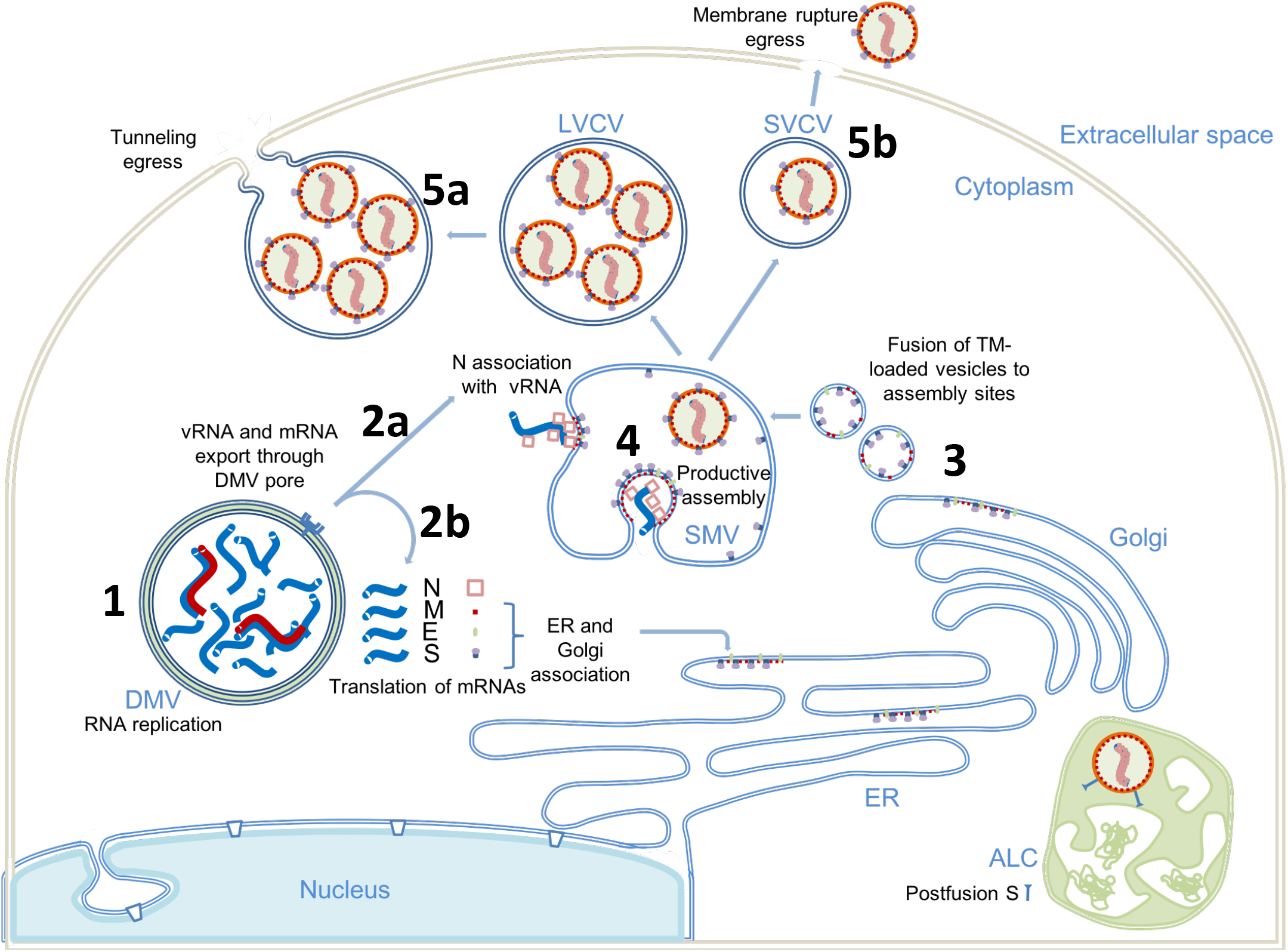
Proposed model of SARS-CoV-2 replication. (1) Viral genome replication occurs inside the DMVs, generating the negative strand viral RNA (red), positive vRNA genomic copy and subgenomic mRNAs (blue). (2) Positive RNAs are exported to cytoplasm through the DMV pores. Subgenomic mRNAs are translated (2b). Structural proteins M, E and S associate with ER, Golgi and ERGIC membranes. Genomic vRNA becomes complexed with newly-synthetized N (2a). (3) S, E and M are transported in dense vesicles which are fuse with the ERGIC SMVs. (4) Productive viral assembly happens in the SMV clustering the viral spikes and encapsidating the genome in RNPs. Viruses bud to the internal space of the SMV. (5) Egress occurs through tunnels via exocytosis-like release (5a) or through membrane rupture (5b). The non-productive autophagolysosome-like compartment (ALC) is depicted in green. DMV, double membrane vesicle; SMV, single membrane vesicle; ALC, autophagolysosome-like compartment; LVCV, large virus-containing vesicle; SVCV, Single virus-containing vesicle.

The genome replication, assembly and egress of the virus is a multistage process that is critically important as it bears the means of medical intervention to stop infection. There are many aspects of this process await further investigation to dissect the mechanism of SARS-CoV-2 replication, including the roles of other viral proteins, such as M and E, as well as host proteins and machines. Nevertheless, this study provides a first glimpse of the SARS-CoV-2 replication cycle under near-native conditions. The methodologies and workflow developed through this study can be broadly applied to studies of infection processes of many other viruses or bacteria, beyond SARS-CoV-2.

## Acknowledgments

We thank Ervin Fodor for helpful discussions, critical reading of manuscript, and facilitating the access to Containment Level 3 lab. We thank Yanan Zhu for help with segmentation and eBIC staff for technical support. We acknowledge Diamond Light Source for access and support of the CryoEM facilities at the UK national electron bio-imaging centre (eBIC, proposals BI26987 and NT21004), funded by the Wellcome Trust, MRC and BBSRC. This research was supported by the National Institutes of Health grant P50AI150481, the UK Wellcome Trust Investigator Award 206422/Z/17/Z, the UK Biotechnology and Biological Sciences Research Council grant BB/S003339/1, and the grant from the Chinese Academy of Medical Sciences Oxford Institute. Containment level 3 experiments were funded through the generous support of philanthropic donors to the University of Oxford’s COVID-19 Research Response Fund. M.L.K. is supported by the Biotechnology and Biological Sciences Research Council (BBSRC) (grant number BB/M011224/1).

## Author contributions

P.Z. conceived the research and with M.H., M.L.M-F., W.J. designed the study. A.H. collected cryoET data. J.B.G. did targeted cryoFIB milling of lamellas. L.C.Z-D., B.B. collected serial cryoFIB/SEM volume imaging data. A.H., L.M., A-S.K., J.R., Y.S., V.D.L. and D.S. performed cryoET 3D reconstructions and analyses. D.S. performed subtomogram averaging. M.L.K. and L.M did viral infections and sample preparation. L.C., L.C.Z-D., D.S. and M.S. did 3D segmentation. I.K., M.A.K., L.M. and M.H. acquired and processed soft X-ray cryo-tomography data. L.M. and P.Z. wrote the manuscript with support from all co-authors.

## Methods

### Sample preparation

Vero Ccl-81 cells (ATCC) were maintained Dulbecco Modified Eagle media supplemented with 5% Fetal Bovine Serum 10 units/mL penicillin (Gibco), 10 μg/mL streptomycin (Gibco), and 2mM l-glutamine. 16,000 cells were seeded on the carbon-side of fibronectin treated G300F1 R2/2 gold EM grids in a 6-well plate well. Infections were performed using passage 3 of SARS-CoV-2 England/02/2020 at MOI of 0.5, 0.1 or 0 (for negative controls). Media was removed from the Vero Ccl-81 cells (ATCC) and replaced with an appropriate amount of virus diluted in 0.5 mL of Dulbecco’s modified Eagle medium (Merck) with 1% FCS, 10 units/mL penicillin (Gibco), 10 μg/mL streptomycin (Gibco), and 2mM l-glutamine. The cells were incubated at room temperature for 15 minutes after which a further 1.5 mL of media was added to each well. The plate was then incubated at 37 °C for 24 hours following which supernatants were discarded and cells washed with 2 mL of PBS. The cells were then fixed by addition of 3 mL of 4% paraformaldehyde in PBS for 1 hour at room temperature. After fixation, grids were plunge-frozen on a Leica Grid plunger 2. 1 ul of concentrated 10 nm gold fiducials was applied to the gold-side of the EM grid and blotted from the gold-side. The grid was quickly immersed in liquid ethane after blotting. Frozen grids were stored in liquid nitrogen until data collection.

### CryoET data acquisition

Tilt series acquisition was carried out at a FEI Titan Krios G2 (Thermo Fisher Scientific) electron microscope operated at 300 kV and equipped with a Gatan BioQuantum energy filter and post-GIF K3 detector (Gatan, Pleasanton, CA).

Tilt series were recorded using SerialEM tilt series controller with pixel sizes of 1.63 Å, 2.13 Å and 4.58 Å for intact cells and 2.13 Å and 7.58 Å on lamella. Zero-loss imaging was used for all tilt series with a 20 eV slit width. Defocus values ranged from −2 μm to −7 μm, except for lamella at 7.58 Å pixel size where 50 μm defocus was used. A 100 μm objective aperture was inserted. A grouped dose-symmetric scheme was used for all tilt-series; intact cells were collected with a range of +/-60 degrees at 3 degree increments in groups of 3 and total dose of 120-135 e/Å^2^; lamella with +/-54 degrees at 3 degree increments and groups of 3 with total dose of 120-135 e/Å^2^ at 4.58 Å and +/-54 degrees at 3 degree increments and groups of 10 with total dose of 70-90 e/ Å^2^ at 7.58 Å. Autofocus and tracking was performed at each tilt with drift measurement taken at tilt reversals with a 10 Å/s target rate. At each tilt, 5 movie frames were recorded using Correlated Double Sampling (CDS) in super-resolution mode and saved in lzw compressed tif format with no gain normalisation. Movies were subsequently gain normalised during motion correction and fourier cropped back to physical pixel size. After each tilt-series a script was run to take a fresh dark reference and reset the defocus offset.

### CryoFIB lamella preparation

Milling of SARS-Cov-2 infected cells was carried out using a Scios DualBeam cryoFIB (ThermoFisher Scientific) equipped with a PP3010T transfer system and stage (Quorum Technologies). Grids were sputter coated within the PP3010T transfer chamber maintained at −175 °C. After loading onto the Scios stage at −168 °C, the grids were inspected using the SEM (operated at 5 kV and 13 pA) and cells, identified as infected from TEM, were found. The grid surface was coated using the gas injection system (Trimethyl(methylcyclopentadienyl)platinum(IV), ThermoFisher Scientific) for 3 s, yielding a thickness of ~3 um. Milling was performed using the ion beam operated at 30 kV and currents decreasing from 300 pA to 30 pA. At 30 pA lamella thickness was less than 300 nm. During the final stage of milling, SEM inspection of the lamellae was conducted at 2 kV and 13 pA.

### Serial cryoFIB/SEM volume imaging

Samples were imaged on a Zeis Crossbeam 550XL fitted with a Quorum transfer station and cryo-stage. They were mounted on a Quorum-compatible custom sample holder and coated with platinum for 60 sec at 10 mA on the Quorum transfer stage, prior to loading on the cryo-stage. Stage temperature was set at −165°C, while the anticontaminator was held at −185°C. Samples were imaged at 45° tilt after being coated again with Pt for 2x 30sec using the FIB-SEM’s internal GIS system, with the Pt reservoir held at 25°C. Initial trapezoid trenches were milled at 30kV 7nA over 15 μm to reach a final depth of 10 μm, with a polish step over a rectangular box with a depth of 10 μm performed at 30kV 1.5 nA. Serial Sectioning and Imaging acquisition was performed as follows: FIB milling was set up using the 30kV 700 pA probe, a z-slice step of 20 nm and a depth of 10 μm over the entire milling box; SEM imaging was performed at a pixel depth of 3024×2304 pixels, which resulted in a pixel size of 6.5 nm, with the beam set at 2kV 35pA, dwell time 100 nsec and scan speed 1, averaging the signal over 100 line scans as a noise-reduction strategy.

### CryoET image processing

The frames in each tilt angle in a tilt series were processed to correct drift using MotionCor2 (Zheng et al., 2017). For the intact cells dataset, all tilt series were aligned using the default parameters in IMOD version 4.10.22 with the eTomo interface, using gold-fiducial markers (Kremer et al., 1996). For lamella dataset, The tilt series were aligned in the framework of Appion-Protomo fiducial-less tilt-series alignment suite (Noble and Stagg, 2015). After tilt series alignment, the tilt-series stacks together with the files describing the projection transformation and fitted tilt angles were transferred to emClarity for the subsequent sub-tomogram averaging analysis (Himes and Zhang, 2018).

### Subtomogram averaging

All sub-tomogram averaging analysis steps were performed using emClarity, mostly following previously published protocols described workflow (Himes and Zhang, 2018). The CTF estimation for each tilt was performed by using emClarity version 1.4.3, and the subvolumes were selected by using automatic template matching function within emClarity using reference derived from EMDB-21452 (Walls et al., 2020) that was low-pass filtered to 30-Å resolution in emClarity. The template matching results were cleaned manually by comparison of the binned tomograms overlaid with the emClarity-generated IMOD model showing the x,y,z coordinates of each cross-correlation peak detected. After manually template cleaning, A total of 450 subvolumes from the lamella dataset and a total of 7090 subvolumes from the extracellular viruses dataset were retained, deriving from 3 tilt-series and 50 tilt-series respectively, for the following averaging and alignment steps in emClarity. For the extracellular viruses dataset, the 3D iterative averaging and alignment procedures were carried out gradually with binning of 4x, 3x, 2x, each with 2-3 iterations with increasingly restrictive search angles and translational shifts. 3-fold symmetry was applied during all the steps. Final converged average map was generated using bin2 tomograms with pixel size of 3.26 Å/pixel and a box size of 123×123×123 voxels. Resolution indicated by 0.143 FSC cut-off was 8.7 Å. The same process was carried out for lamella dataset, except for the final average map was generated with pixel size of 4.26 Å/pixel and a box size of 90×90×90 voxels and a final resolution at 11Å (Gold standard FSC at 0.143 cut-off).

### Serial cryoFIB/SEM Segmentation

Cell structures were manually segmented from stacks of images using ImageJ (Koppensteiner et al., 2012) and Microscopy Image Browser (MIB) software (Belevich et al., 2016) on a Windows computer with 32GB RAM and Wacom Cintiq Pro display tablet with pen. Datasets of below 2GB in .mrc format were analysed one at a time, where one dataset comprised of 200 subsequent images on average.

### CryoET segmentation and 3D visualization

Transport vesicles, Viral membrane, Nuclear membrane, Double membrane vesicles (DMV), and single membrane vesicles (SMV) were segmented using Convolutional Neural Networks based tomogram annotation in the EMAN2.2 software package (Chen et al., 2017). Viral spikes were mapped back to their original particles position using emClarity tomoCPR function. UCSF Chimera (Pettersen et al., 2004) was used to visualize the segmentations and subtomogram average structures in 3D.

### Soft X-ray Cryo-tomography

Data were collected in areas of interest on vitrified samples on 3mm TEM grids according to established protocols (Kounatidis et al., 2020). Grids were loaded on the X-ray microscope at B24 and were first mapped using visible light with a 20X objective. The resulting coordinate-map was used to locate areas of interest where 2D X-ray mosaics were collected (X-ray light used was at 500eV) and used to identify areas of interest within. Tilt series of 100-120 ° were collected for each field of view area of interest at 0.2 or 0.5° steps with constant exposure of 0.5 sec keeping average pixel intensity to between 5-30k counts. All tilt series were background subtracted, saved as raw Tiff stacks and reconstructed using either IMOD (Kremer et al., 1996) or Batchruntomo (Mastronarde, 2005).

### Quantification and statistical analyses

Number of pores in DMV and plasma membrane discontinuities were determined after visual inspection and visual counting by two independent investigators. The investigators were not blinded to allocation during experiments and outcome assessment.

## References

Andrejeva, J., Childs, K.S., Young, D.F., Carlos, T.S., Stock, N., Goodbourn, S., and Randall, R.E. (2004). The V proteins of paramyxoviruses bind the IFN-inducible RNA helicase, mda-5, and inhibit its activation of the IFN-beta promoter. Proc Natl Acad Sci U S A 101, 17264–17269.

Belevich, I., Joensuu, M., Kumar, D., Vihinen, H., and Jokitalo, E. (2016). Microscopy Image Browser: A Platform for Segmentation and Analysis of Multidimensional Datasets. PLoS Biol 14, e1002340.

Belouzard, S., Chu, V.C., and Whittaker, G.R. (2009). Activation of the SARS coronavirus spike protein via sequential proteolytic cleavage at two distinct sites. Proceedings of the National Academy of Sciences of the United States of America 106, 5871–5876.

Berger, W., Steiner, E., Grusch, M., Elbling, L., and Micksche, M. (2009). Vaults and the major vault protein: Novel roles in signal pathway regulation and immunity. Cellular and Molecular Life Sciences 66, 43–61.

Cai, Y., Zhang, J., Xiao, T., Peng, H., Sterling, S.M., Walsh, R.M., Rawson, S., Rits-Volloch, S., and Chen, B. (2020). Distinct conformational states of SARS-CoV-2 spike protein. Science 1592, eabd4251–eabd4251.

Chen, M., Dai, W., Sun, S.Y., Jonasch, D., He, C.Y., Schmid, M.F., Chiu, W., and Ludtke, S.J. (2017). Convolutional neural networks for automated annotation of cellular cryo-electron tomograms. Nat Methods 14, 983–985.

de Wilde, A.H., Raj, V.S., Oudshoorn, D., Bestebroer, T.M., van Nieuwkoop, S., Limpens, R.W.A.L., Posthuma, C.C., van der Meer, Y., Bárcena, M., Haagmans, B.L., et al. (2013). MERS-coronavirus replication induces severe in vitro cytopathology and is strongly inhibited by cyclosporin A or interferon-α treatment. Journal of General Virology 94, 1749–1760.

Doyle, N., Neuman, B.W., Simpson, J., Hawes, P.C., Mantell, J., Verkade, P., Alrashedi, H., and Maier, H.J. (2018). Infectious bronchitis virus nonstructural protein 4 alone induces membrane pairing. Viruses 10, 1–17.

Ertel, K.J., Benefield, D., Castaño-Diez, D., Pennington, J.G., Horswill, M., Den Boon, J.A., Otegui, M.S., and Ahlquist, P. (2017). Cryo-electron tomography reveals novel features of a viral rna replication compartment. eLife 6, 1–24.

Fan, X., Cao, D., Kong, L., and Zhang, X. (2020). Cryo-EM analysis of the post-fusion structure of the SARS-CoV spike glycoprotein. Nature Communications 11, 1–10.

Fried, J.A., Ramasubbu, K., Bhatt, R., Topkara, V.K., Clerkin, K.J., Horn, E., Rabbani, L.R., Brodie, D., Jain, S.S., Kirtane, A.J., et al. (2020). The variety of cardiovascular presentations of COVID-19. Circulation, 1930–1936.

Goldsmith, C.S., Tatti, K.M., Ksiazek, T.G., Rollin, P.E., Comer, J.A., Lee, W.W., Rota, P.A., Bankamp, B., Bellini, W.J., and Zaki, S.R. (2004). Ultrastructural Characterization of SARS Coronavirus. Emerging Infectious Diseases 10, 320–326.

Hagemeijer, M.C., Monastyrska, I., Griffith, J., van der Sluijs, P., Voortman, J., van Bergen en Henegouwen, P.M., Vonk, A.M., Rottier, P.J.M., Reggiori, F., and De Haan, C.A.M. (2014). Membrane rearrangements mediated by coronavirus nonstructural proteins 3 and 4. Virology 458-459, 125–135.

Harkiolaki, M., Darrow, M.C., Spink, M.C., Kosior, E., Dent, K., and Duke, E. (2018). Cryo-soft X-ray tomography: using soft X-rays to explore the ultrastructure of whole cells. Emerging Topics in Life Sciences 2, 81–92.

Himes, B.A., and Zhang, P. (2018). emClarity: software for high-resolution cryo-electron tomography and subtomogram averaging. Nat Methods 15, 955961.

Hoffmann, M., Kleine-Weber, H., Schroeder, S., Krüger, N., Herrler, T., Erichsen, S., Schiergens, T.S., Herrler, G., Wu, N.H., Nitsche, A., et al. (2020). SARS-CoV-2 Cell Entry Depends on ACE2 and TMPRSS2 and Is Blocked by a Clinically Proven Protease Inhibitor. Cell 181, 271–280.e278.

Hornung, V., Ellegast, J., Kim, S., Brzozka, K., Jung, A., Kato, H., Poeck, H., Akira, S., Conzelmann, K.K., Schlee, M., et al. (2006). 5’-Triphosphate RNA is the ligand for RIG-I. Science 314, 994–997.

Ke, Z., Oton, J., Qu, K., Cortese, M., Zila, V., McKeane, L., Nakane, T., Zivanov, J., Neufeldt, C.J., Cerikan, B., et al. (2020). Structures and distributions of SARS-CoV-2 spike proteins on intact virions. Nature.

Knoops, K., Kikkert, M., Van Den Worm, S.H.E., Zevenhoven-Dobbe, J.C., Van Der Meer, Y., Koster, A.J., Mommaas, A.M., and Snijder, E.J. (2008). SARS-coronavirus replication is supported by a reticulovesicular network of modified endoplasmic reticulum. PLoS Biology 6, 1957–1974.

Koppensteiner, H., Banning, C., Schneider, C., Hohenberg, H., and Schindler, M. (2012). Macrophage internal HIV-1 is protected from neutralizing antibodies. J Virol 86, 2826–2836.

Kounatidis, I., Stanifer, M.L., Phillips, M.A., Paul-Gilloteaux, P., Heiligenstein, X., Wang, H., Okolo, C.A., Fish, T.M., Spink, M.C., Stuart, D.I., et al. (2020). 3D Correlative Cryo-Structured Illumination Fluorescence and Soft X-ray Microscopy Elucidates Reovirus Intracellular Release Pathway. Cell 182, 515–530.e517.

Kremer, J.R., Mastronarde, D.N., and McIntosh, J.R. (1996). Computer visualization of three-dimensional image data using IMOD. J Struct Biol 116, 71–76.

Lamers, M.M., Beumer, J., Vaart, J.V.D., Knoops, K., Puschhof, J., Breugem, T.I., Ravelli, R.B.G., Schayck, J.P.V., Mykytyn, A.Z., Duimel, H.Q., et al. (2020). SARS-CoV-2 productively infects human gut enterocytes. Science 369, 50–54.

Lan, J., Ge, J., Yu, J., Shan, S., Zhou, H., Fan, S., Zhang, Q., Shi, X., Wang, Q., Zhang, L., et al. (2020). Structure of the SARS-CoV-2 spike receptor-binding domain bound to the ACE2 receptor. Nature 581, 215–220.

Lu, S., Ye, Q., Singh, D., Villa, E., Cleveland, D.W., and Corbett, K.D. (2020). The SARS-CoV-2 Nucleocapsid phosphoprotein forms mutually exclusive condensates with RNA and the membrane-associated M protein. bioRxiv: the preprint server for biology.

Lucas, C., Wong, P., Klein, J., Castro, T.B.R., Silva, J., Sundaram, M., Ellingson, M.K., Mao, T., Oh, J.E., Israelow, B., et al. (2020). Longitudinal analyses reveal immunological misfiring in severe COVID-19. Nature 584.

Mastronarde, D.N. (2005). Automated electron microscope tomography using robust prediction of specimen movements. J Struct Biol 152, 36–51.

Neufeldt, C.J., Joyce, M.A., Van Buuren, N., Levin, A., Kirkegaard, K., Gale, M., Tyrrell, D.L.J., and Wozniak, R.W. (2016). The Hepatitis C Virus-Induced Membranous Web and Associated Nuclear Transport Machinery Limit Access of Pattern Recognition Receptors to Viral Replication Sites. PLoS Pathogens 12, 1–28.

Neuman, B.W., Kiss, G., Kunding, A.H., Bhella, D., Baksh, M.F., Connelly, S., Droese, B., Klaus, J.P., Makino, S., Sawicki, S.G., et al. (2011). A structural analysis of M protein in coronavirus assembly and morphology. Journal of Structural Biology 174, 11–22.

Noble, A.J., and Stagg, S.M. (2015). Automated batch fiducial-less tilt-series alignment in Appion using Protomo. Journal of Structural Biology 192, 270–278.

Ogando, N.S., Dalebout, T.J., Zevenhoven-Dobbe, J.C., Limpens, R.W.A.L., van der Meer, Y., Caly, L., Druce, J., de Vries, J.J.C., Kikkert, M., Bárcena, M., et al. (2020). SARS-coronavirus-2 replication in Vero E6 cells: replication kinetics, rapid adaptation and cytopathology. Journal of General Virology.

Oudshoorn, D., Rijs, K., Limpens, R.W.A.L., Groen, K., Koster, A.J., Snijder, E.J., Kikkert, M., and Bárcena, M. (2017). Expression and cleavage of middle east respiratory syndrome coronavirus nsp3-4 polyprotein induce the formation of double-membrane vesicles that mimic those associated with coronaviral RNA replication. mBio 8, 1–17.

Paul, D., Hoppe, S., Saher, G., Krijnse-Locker, J., and Bartenschlager, R. (2013). Morphological and Biochemical Characterization of the Membranous Hepatitis C Virus Replication Compartment. Journal of Virology 87, 10612–10627.

Pettersen, E.F., Goddard, T.D., Huang, C.C., Couch, G.S., Greenblatt, D.M., Meng, E.C., and Ferrin, T.E. (2004). UCSF Chimera--a visualization system for exploratory research and analysis. J Comput Chem 25, 1605–1612.

Reggiori, F., Monastyrska, I., Verheije, M.H., Calì, T., Ulasli, M., Bianchi, S., Bernasconi, R., De Haan, C.A.M., and Molinari, M. (2010). Coronaviruses hijack the LC3-I-positive EDEMosomes, ER-derived vesicles exporting short-lived ERAD regulators, for replication. Cell Host and Microbe 7, 500–508.

Shang, J., Ye, G., Shi, K., Wan, Y., Luo, C., Aihara, H., Geng, Q., Auerbach, A., and Li, F. (2020). Structural basis of receptor recognition by SARS-CoV-2. Nature 581, 221–224.

Sharma, A., Garcia, G., Wang, Y., Plummer, J.T., Morizono, K., Arumugaswami, V., and Svendsen, C.N. (2020). Human iPSC-Derived Cardiomyocytes Are Susceptible to SARS-CoV-2 Infection. Cell Reports Medicine 1, 100052–100052.

Simmons, G., Reeves, J.D., Rennekamp, A.J., Amberg, S.M., Piefer, A.J., and Bates, P. (2004). Characterization of severe acute respiratory syndrome-associated coronavirus (SARS-CoV) spike glycoprotein-mediated viral entry. Proceedings of the National Academy of Sciences of the United States of America 101, 4240–4245.

Snijder, E.J., Limpens, R.W.A.L., de Wilde, A.H., de Jong, A.W.M., Zevenhoven-Dobbe, J.C., Maier, H.J., Faas, F.F.G.A., Koster, A.J., and Bárcena, M. (2020). A unifying structural and functional model of the coronavirus replication organelle: Tracking down RNA synthesis. PLoS Biology 18, 1–25.

Song, W., Gui, M., Wang, X., and Xiang, Y. (2018). Cryo-EM structure of the SARS coronavirus spike glycoprotein in complex with its host cell receptor ACE2. 14, e1007236.

Stertz, S., Reichelt, M., Spiegel, M., Kuri, T., Martínez-Sobrido, L., García-Sastre, A., Weber, F., and Kochs, G. (2007). The intracellular sites of early replication and budding of SARS-coronavirus. Virology 361, 304315.

Surya, W., Li, Y., and Torres, J. (2018). Structural model of the SARS coronavirus E channel in LMPG micelles. Biochimica et Biophysica Acta - Biomembranes 1860, 1309–1317.

Sutton, G., Sun, D., Fu, X., Kotecha, A., Hecksel, C.W., Clare, D.K., Zhang, P., Stuart, D.I., and Boyce, M. (2020). Assembly intermediates of orthoreovirus captured in the cell. Nature Communications 11, 1–7.

Toelzer, C., Gupta, K., Yadav, S.K.N., Borucu, U., Davidson, A.D., Kavanagh Williamson, M., Shoemark, D.K., Garzoni, F., Staufer, O., Milligan, R., et al. (2020). Free fatty acid binding pocket in the locked structure of SARS-CoV-2 spike protein. Science.

Turoňová, B., Sikora, M., Schürmann, C., Hagen, W., Welsch, S., Blanc, F., von Bülow, S., Gecht, M., Bagola, K., Hörner, C., et al. (2020). In situ structural analysis of SARS-CoV-2 spike reveals flexibility mediated by three hinges. 5223, 1–12.

Vennema, H., Godeke, G.J., Rossen, J.W.A., Voorhout, W.F., Horzinek, M.C., Opstelten, D.J.E., and Rottier, P.J.M. (1996). Nucleocapsid-independent assembly of coronavirus-like particles by co-expression of viral envelope protein genes. EMBO Journal 15, 2020–2028.

Walls, A.C., Park, Y.J., Tortorici, M.A., Wall, A., McGuire, A.T., and Veesler, D. (2020). Structure, Function, and Antigenicity of the SARS-CoV-2 Spike Glycoprotein. Cell 181, 281–292.e286.

Wang, Q., Zhang, Y., Wu, L., Niu, S., Song, C., Zhang, Z., Lu, G., Qiao, C., Hu, Y., Yuen, K.Y., et al. (2020). Structural and Functional Basis of SARS-CoV-2 Entry by Using Human ACE2. Cell 181, 894–904.e899.

Wolff, G., Limpens, R.W.A.L., Zevenhoven-Dobbe, J.C., Laugks, U., Zheng, S., de Jong, A.W.M., Koning, R.I., Agard, D.A., Grünewald, K., Koster, A.J., et al. (2020a). A molecular pore spans the double membrane of the coronavirus replication organelle. Science 3629, eabd3629–eabd3629.

Wolff, G., Melia, C.E., Snijder, E.J., and Bárcena, M. (2020b). Double–Membrane Vesicles as Platforms for Viral Replication. Trends in Microbiology.

Woodward, C.L., Mendonca, L.M., and Jensen, G.J. (2015). Direct visualization of vaults within intact cells by electron cryo-tomography. Cell Mol Life Sci 72, 3401–3409.

Yan, R., Zhang, Y., Li, Y., Xia, L., Guo, Y., and Zhou, Q. (2020). Structural basis for the recognition of the SARS-CoV-2 by full-length human ACE2. Science.

Yao, H., Song, Y., Chen, Y., Wu, N., Xu, J., Sun, C., Zhang, J., Weng, T., Zhang, Z., Wu, Z., et al. (2020). Molecular architecture of the SARS-CoV-2 virus.

Zheng, S.Q., Palovcak, E., Armache, J.P., Verba, K.A., Cheng, Y., and Agard, D.A. (2017). MotionCor2: anisotropic correction of beam-induced motion for improved cryo-electron microscopy. Nat Methods 14, 331–332.

Zhou, X., Cong, Y., Veenendaal, T., Klumperman, J., Shi, D., Mari, M., and Reggiori, F. (2017). Ultrastructural characterization of membrane rearrangements induced by porcine epidemic diarrhea virus infection. Viruses 9.

Zhu, Y., Sun, D., Schertel, A., Martin-fernandez, M.L., Freyberg, Z., Zhang, P., Ning, J., Fu, X., Gwo, P.P., and Watson, A.M. (2021). Short Article Serial cryoFIB / SEM Reveals Cytoarchitectural Disruptions in Leigh Syndrome Patient Cells Serial cryoFIB / SEM Reveals Cytoarchitectural Disruptions in Leigh Syndrome Patient Cells. Structure/Folding and Design, 1–6.

