## Supplemental figures for "SARS-CoV-2 Assembly and Egress Pathway Revealed by Correlative Multi-modal Multi-scale Cryo-imaging"

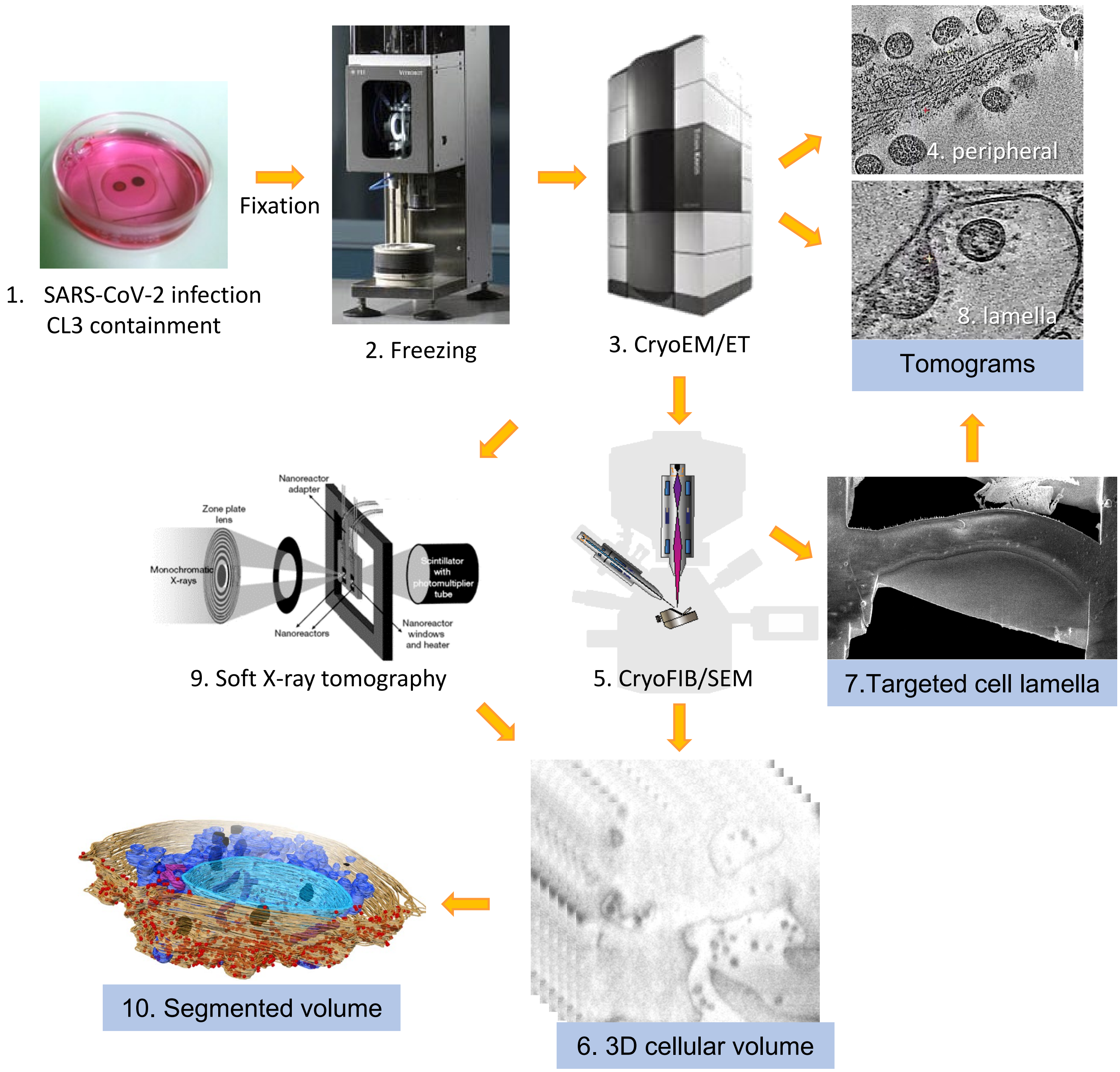


**Figure S1 | A workflow for correlative multimodal multiscale imaging of SARS-CoV-2 infected cells.** 1) Cells are grown on indexed EM grids, infected with SARS-CoV-2 and fixed with paraformaldehyde. 2) Grids are plunge-frozen in liquid ethane and 3) imaged by cryoEM/ET to locate the infected cells. 4) Tomograms are collected on the cell periphery of infected cells. 5) Infected cells are subjected to processing and imaging in a cryoFIB/SEM dualbeam instrument for 6) serial cryoFIB/SEM volume imaging and 7) targeted cell lamella. 8) Tomograms are collected from cell lamellae. 9) Alternatively, infected cells are imaged by soft-X-ray cryo-tomography. 10) Cellular volume data are manually segmented.

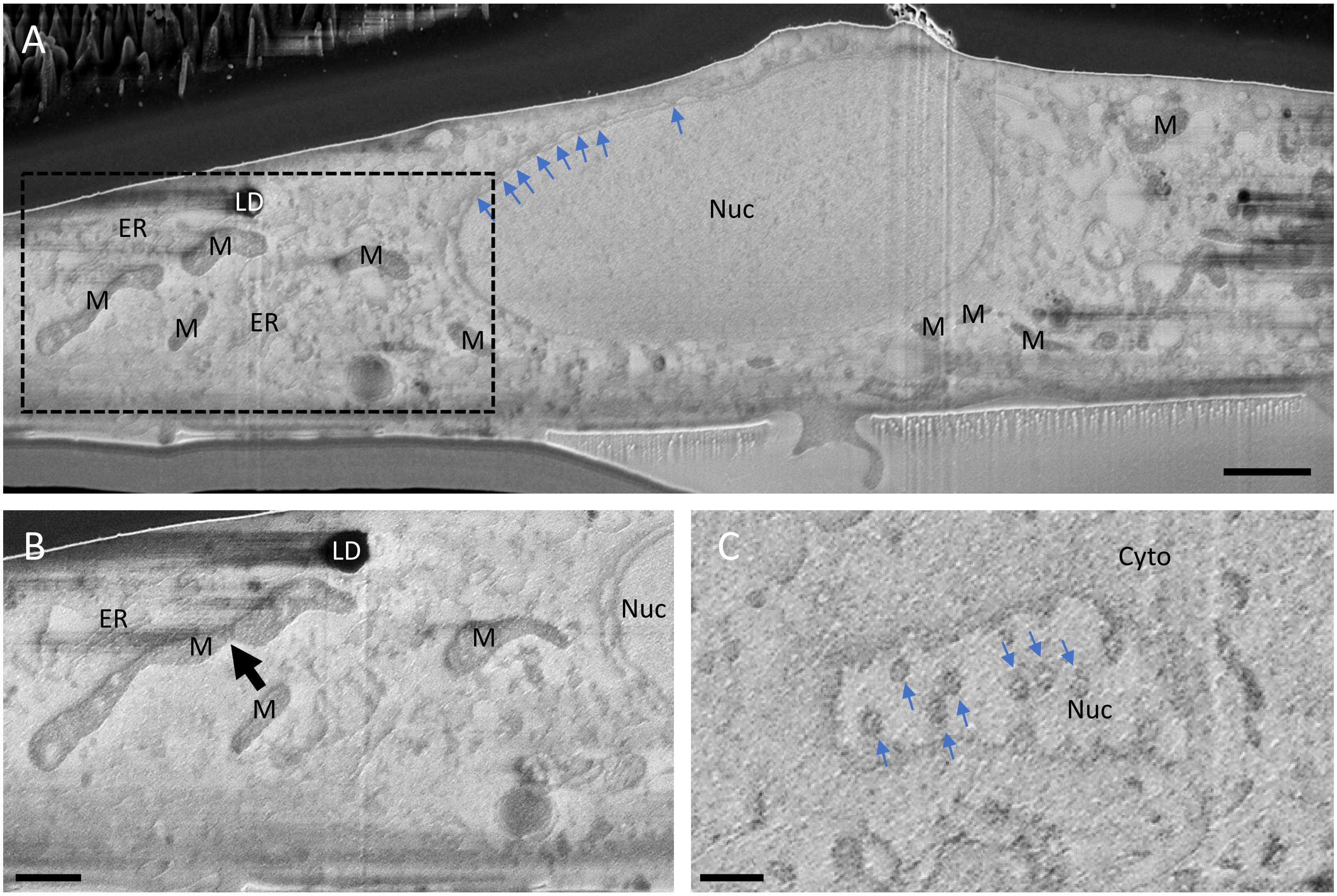


**Figure S2 | Serial cryoFIB/SEM of control uninfected cell.** (A) A representative overview of a cryoFIB/SEM slice from an uninfected cell. Blue arrows point to nuclear pores. (B) Detailed view of the dashed area in A from a slice 100 nm in depth depicting a connected mitochondrial network. (C) A cryoFIB/SEM slice tangential to the nuclear envelope, showing top view of nucleopores (blue arrows). Nuc, nucleus; Cyto, cytoplasm; ER, endoplasmic reticulum; M, mitochondria; LD, lipid droplet. Scale bar is 600 nm in A, 300 nm in B and C.

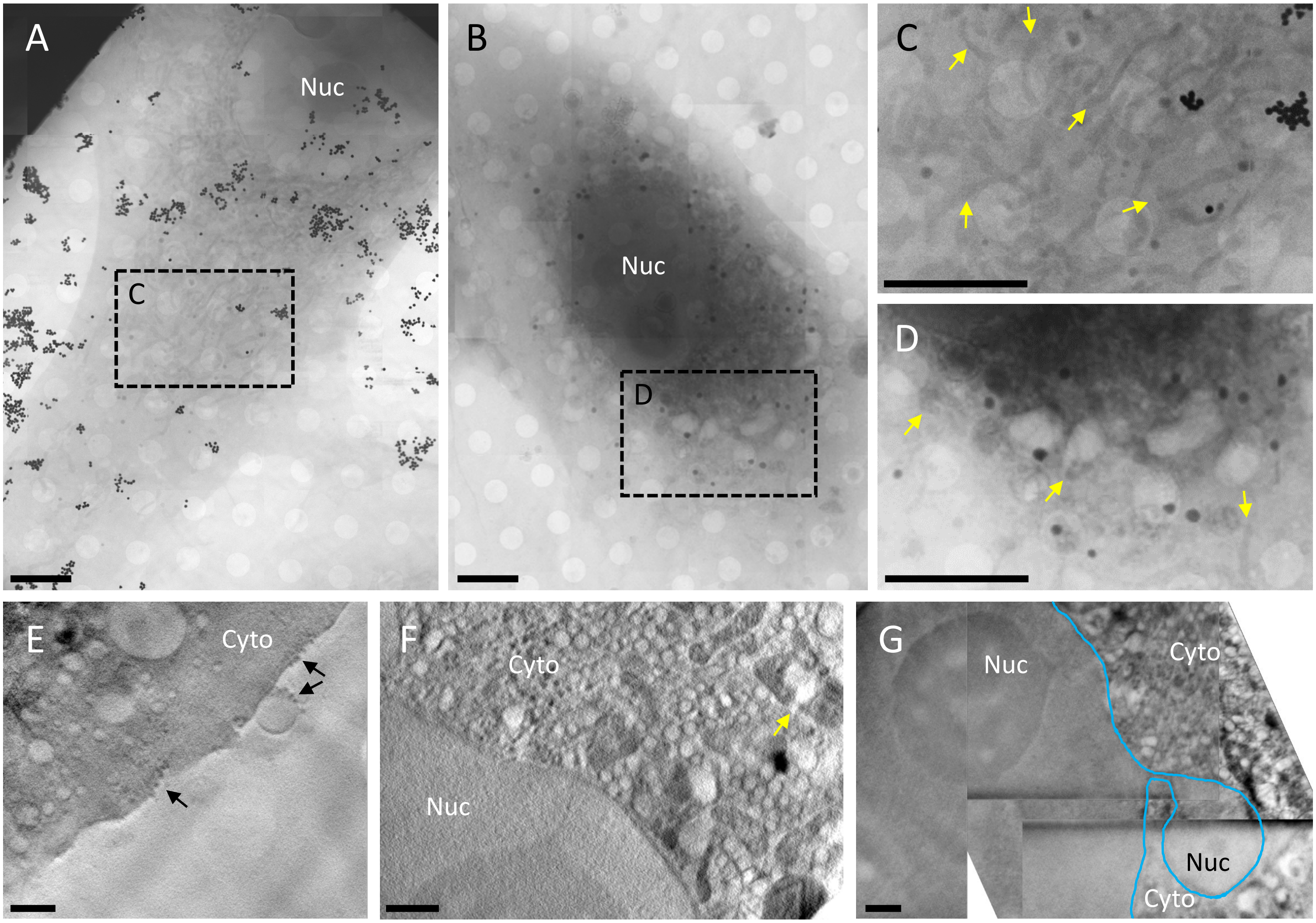


**Figure S3 | Soft X-ray cryo-tomography of SARS-CoV-2 infected cells.** (A-B) Soft X-ray overview mosaics of uninfected (A) and infected (B) Vero cells. (C-D) Detailed view of boxed area in A and B depicting mitochondrial network in uninfected (C) and fragmented mitochondria in infected cell (D) (yellow arrows). (E-F) Soft X-ray tomogram slices taken from infected cells depicting viruses at cell edge (E) (black arrows), abundant DMVs and a damaged mitochondria (F) (yellow arrow). (G) A montage of four tomograms depicting cytoplasmic invasion (or nuclear blebbing) in an infected cell. Nuclear envelope is outlined in cyan. Nuc, nucleus; Cyto, cytoplasm. Black dots in A and C are gold fiducial markers. Scale bar is 5 µm in A, B, C, D; 1 µm in E, F and G.

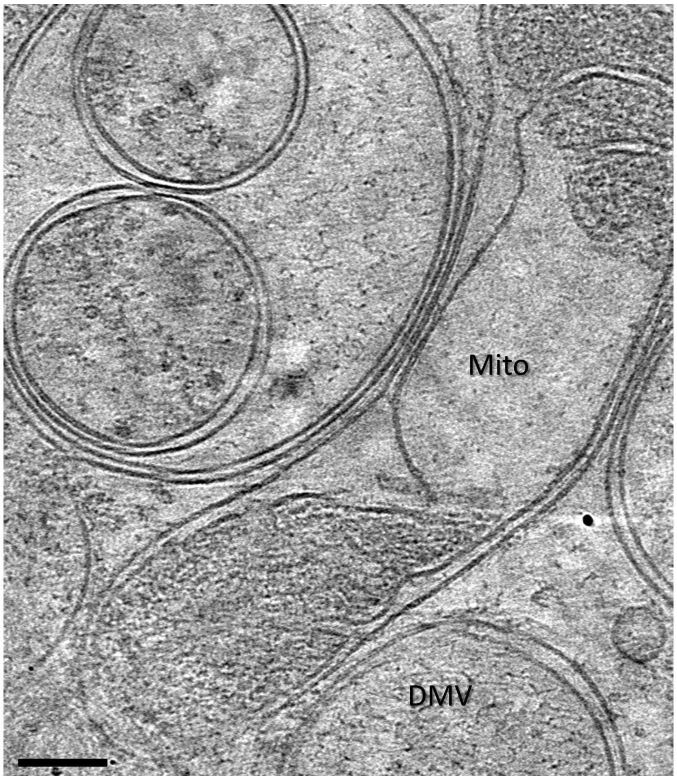


**Figure S4 | Effect of SARS-CoV-2 replication on mitochondria**. Representative damaged mitochondria in SARS-CoV-2 infected cells as seen in a cryoET slice from a cell lamella. Mitochondria and DMV are labeled.

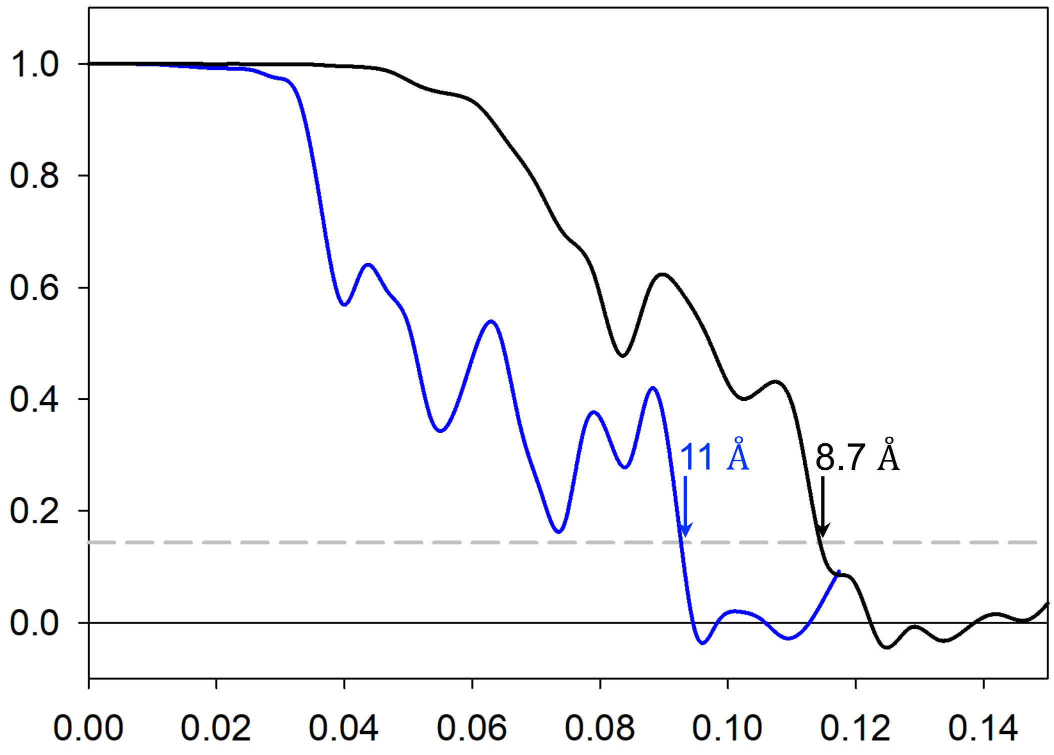


**Figure S5 |** Fourier shell correlation (FSC) plots of subtomogram averaged spike density maps derived from intracellular virions (blue, from 450 subvolumes) and from released virions (black, from 7090 subvolumes). The dashed lines mark the FSC value of 0.143.

Movie 1. Serial cryoFIB/SEM of an uninfected cell
Movie 2. Serial cryoFIB/SEM of an infected cell
Movie 3. Segmentation of the volume depicted in Movie 2.
Movie 4. Serial cryoFIB/SEM of an infected cell, showing abound heterogeneous vesicles
Movie 5. Serial cryoFIB/SEM of an infected cell, showing cytoplasmic invagination
Movie 6. CryoET of DMVs and portals
Movie 7. CryoET of virus assembly sites
Movie 8. CryoET of released viruses outside of a cell
Movie 9. CryoET of virus egress site, broken membrane
